# Pre-exposure to stress reduces loss of community and genetic diversity following severe environmental disturbance

**DOI:** 10.1101/2024.10.11.617827

**Authors:** Charles C.Y. Xu, Vincent Fugère, Naíla Barbosa da Costa, Beatrix E. Beisner, Graham Bell, Melania E. Cristescu, Gregor F. Fussmann, Andrew Gonzalez, B. Jesse Shapiro, Rowan D.H. Barrett

## Abstract

Environmental stress caused by anthropogenic impacts is increasing worldwide. Understanding the ecological and evolutionary consequences for biodiversity will be crucial for our ability to respond effectively. Historical exposure to environmental stress is expected to select for resistant species, shifting community composition towards more stress-tolerant taxa. Concurrent with this species sorting process, genotypes within resistant taxa that have the highest relative fitness under severe stress are expected to increase in frequency, leading to evolutionary adaptation. However, empirical demonstrations of these dual ecological and evolutionary processes in natural communities are lacking. Here, we provide the first evidence for simultaneous species sorting and evolutionary adaptation across multiple species within a natural freshwater bacterial community. Using a two-phase stressor experimental design (acidification pre-exposure followed by severe acidification) in aquatic mesocosms, we show that pre-exposed communities were more resistant than naïve communities to taxonomic loss when faced with severe acid stress. However, after sustained severe acidification, taxonomic richness of both pre-exposed and naïve communities eventually converged. All communities experiencing severe acidification became dominated by an acidophilic bacterium, *Acidiphilium rubrum*, but this species retained greater genetic diversity and followed distinct evolutionary trajectories in pre-exposed relative to naïve communities. These patterns were shared across other acidophilic species, providing repeated evidence for the impact of pre-exposure on evolutionary outcomes despite the convergence of community profiles. Our results underscore the need to consider both ecological and evolutionary processes to accurately predict the responses of natural communities to environmental change.

## Introduction

Any environmental event that reduces individual fitness is defined as a stress, and such stress is common in ecological communities ^1^. Environmental stress is expected to intensify worldwide due to increasing anthropogenic impacts from multiple stressors ^2^. Severe stress can induce widespread mortality, causing significant biodiversity decline and large shifts in community composition and functioning ^3^. Community responses to stress are characterized by resistance (the capacity of a system to withstand disturbance) and resilience (the ability to recover after disturbance) ^4,5^. If community resistance and resilience are insufficient and stress is not alleviated, the biodiversity of a community may collapse. Elucidating the ecological and evolutionary mechanisms preventing or permitting community persistence and recovery after severe environmental stress is crucial for understanding biodiversity change and improving its management and conservation.

Pre-exposure to stress can mediate the response of complex communities via phenotypic plasticity, species sorting, and evolutionary adaptation ^6^. Phenotypic plasticity may enable individual organisms to exhibit induced tolerance after exposure to sublethal levels of stress ^7^. Historical stress may select against susceptible species during pre-exposure, shifting community composition towards more stress-tolerant taxa. This species sorting process can maintain essential ecological interactions and functioning, and prevent community collapse when confronted with levels of stress that would otherwise be lethal in the absence of pre-exposure ^8,9^. While phenotypic plasticity and species sorting are well-known outcomes of pre-exposure to stress ^10,11^, much less is known about the role of evolution in this process, especially in complex natural communities. Historical exposure to environmental stress is expected to select for genotypes with higher relative fitness under severe stress, provided that adaptive alleles are present as standing variation within a species or appear via mutation and have sufficiently large selection coefficients to counter the effects of drift ^12–14^. If an increase in frequency of adaptive genotypes rescues average population fitness, this can ameliorate, prevent, or reverse population decline in response to stress ^15,16^, thereby allowing greater mutational input to maintain levels of genetic diversity ^17,18^. In an analogy to the way that vaccination with a sub-lethal dose of a virus can prepare an individual’s immune system for a potentially deadly future infection, pre-exposure to low or moderate levels of stressors can protect ecological communities from future severe stress. Pre-exposure has been demonstrated to mitigate the impact of severe environmental stress on community structure in a variety of systems ^19–22^, even when the specific stressors are different in each exposure period ^23^, and the effects can extend to community and ecosystem functioning as well ^24–26^.

Acidification is well-known to be a major environmental stressor for aquatic ecosystems, and its detrimental effects on biodiversity have been a considerable challenge for management and conservation ^26–30^. Several whole-ecosystem studies have been conducted on acidification of freshwater lakes, revealing declines in species diversity and disruptions to primary production and nutrient cycling ^31–33^. Even though freshwater acidification continues to be a relevant issue to ecological and human health ^34,35^, understanding of the evolutionary processes underlying community responses in aquatic ecosystems remains limited. This holds for microbial communities despite their capacity for rapid evolution ^36,37^. It is difficult to disentangle evolutionary adaptation from ecological species sorting because their effects can be indistinguishable using standard community profiling techniques such as 16S or 18S rRNA gene metabarcoding. As such, investigating the relative contributions of ecological and evolutionary responses of natural communities requires careful experimentation and monitoring of taxonomic and genetic diversity dynamics over time.

To address this gap, we experimentally manipulated aquatic mesocosms containing diverse microbial communities derived from a natural unpolluted lake to test the impact of acidification pre-exposure on community-wide taxonomic composition and intraspecific genetic diversity after severe acidification. In addition, we tested the role of low-dispersal rate in moderating these effects. Under environmental stress, immigration via dispersal can buffer against demographic decline and help maintain standing genetic and taxonomic variation, but introduction of maladapted types can also decrease average fitness depending on dispersal rates ^8,22,38^. We used 16S metabarcoding to profile microbial communities and metagenomic shotgun sequencing to investigate the potential role of evolutionary adaptation within individual species during community response. We hypothesized that (1) pre-exposure to moderate acidification will select for acidophiles, thereby producing a more resistant community during severe and sustained acidification than naïve communities whose first experience with acidification occurs at severe levels, (2) dispersal will mitigate loss in taxonomic diversity, and (3) species that survive severe acidification in pre-exposed communities will evolve along distinct trajectories compared to those in naïve communities, leading to genetic differentiation and greater resistance to genetic diversity loss. Our experiment provides the first evidence for simultaneous species sorting and evolutionary adaptation across multiple species within a natural community. Broadly, this study provides novel insight into the ecological and evolutionary dynamics of complex microbial communities as they respond to severe environmental stress.

## Results

### Changes in community diversity following severe acidification

We first tested the hypothesis that the ecological response (species sorting) to acid stress would depend on pre-exposure to a milder acid stress. Briefly, the experiment consisted of two phases (Figure 1). In phase I, pre-exposed mesocosms (red) were subjected to weekly acidification treatments of pH 4 while naïve mesocosms (blue) naturally fluctuated around pH 8.5 for approximately seven weeks. In phase II, all mesocosms were subjected to a press acidification treatment of pH 3 for approximately eight weeks apart from four additional control mesocosms (green). Alpha diversity as measured by the Shannon index differed significantly between pre-exposed and naïve communities indicating the effect of pre-exposure on amplicon sequence variant (ASV) richness and evenness (Figure 2A). The experiment began with all mesocosms at approximately a Shannon index of 5.6, but pre-exposure caused a significant decrease by the end of the pre-exposure treatment (Sample 2) (H’ = 18.64, *Q* < 0.05, Benjamini-Hochberg *Q*-value based on *P*-values from Kruskal-Wallis tests) (Figure S1). Although both pre-exposed and naïve communities decreased in diversity one week after severe acidification, naïve communities experienced a significantly greater decline, resulting in lower Shannon values than pre-exposed communities (Sample 3) (H’ = 22.91, *Q* < 0.05). Pre-exposed communities continued to decrease in diversity during the eight weeks of severe acidification while naïve communities recovered slightly such that all treated communities converged on a Shannon index of 1.8 regardless of pre-exposure (Sample 4) (H’ = 1.37, *Q* > 0.05). The diversity of control communities remained unchanged throughout the experiment (Sample 2 – Sample 3: H’ = 2.08, *Q* > 0.05; Sample 3 – Sample 4: H’ = 0.33, *Q* > 0.05). The observed recovery in Shannon diversity of naïve communities by the end of the experiment coincided with a small but significant increase in the number of observed ASVs (Sample 3 – Sample 4: H’ = 17.95, *Q* < 0.05) whereas no significant change was observed in pre-exposed communities (Sample 3 – Sample 4: H’ = 4.11, *Q* > 0.05) (Figure S2). The pairwise change in diversity of pre-exposed communities was significantly negative between all time points whereas in naïve communities it was only significantly negative immediately after severe acidification (Sample 2 – Sample 3) and by the end of the experiment it was significantly positive (Sample 3 – Sample 4), indicating recovery (Table S1; Figure S3). Pairwise change in the Shannon index was significantly different between pre-exposed and naïve communities across all time points (Table S2). Dispersal did not significantly affect Shannon values throughout the experiment (Table S1).

**Figure 1.**
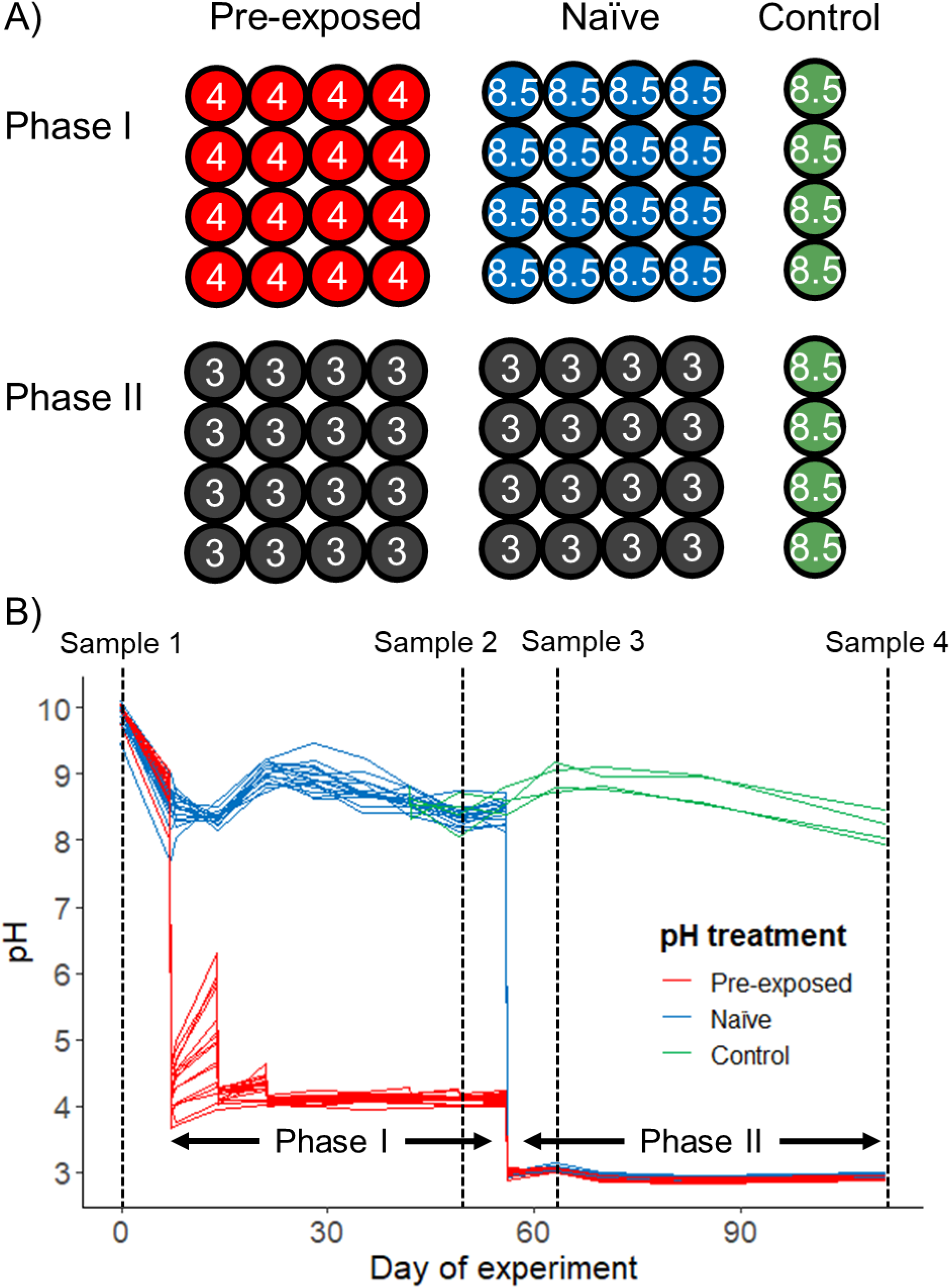
Experimental design and pre-exposure treatments. A) Schematic representation of the subset of mesocosms (circles) from the two-phase experiment included in this study. Colors indicate pre-exposed (red), naïve (blue), and control (green) mesocosms, and numbers indicate pH. Half of pre-exposed and naïve mesocosms were under a global dispersal regime in phase I while the other half were isolated (dispersal regimes not shown). B) Measured pH of each mesocosm throughout the experiment. Each line represents an individual mesocosm, colors indicate pH treatments, and arrows indicate the duration of each experimental phase. Pre-exposed mesocosms were acidified to pH 4 at the beginning of phase I on June 14/day 7, and, in phase II, all mesocosms were acidified to pH 3 on August 2/day 56 until the end of the experiment. Black dashed lines mark the four time points during the experiment when samples were taken (June 7/day 0, July 26/day 49, August 9/day 63, and September 25/day 111) referred to as Sample 1, Sample 2, Sample 3, and Sample 4.

**Figure 2.**
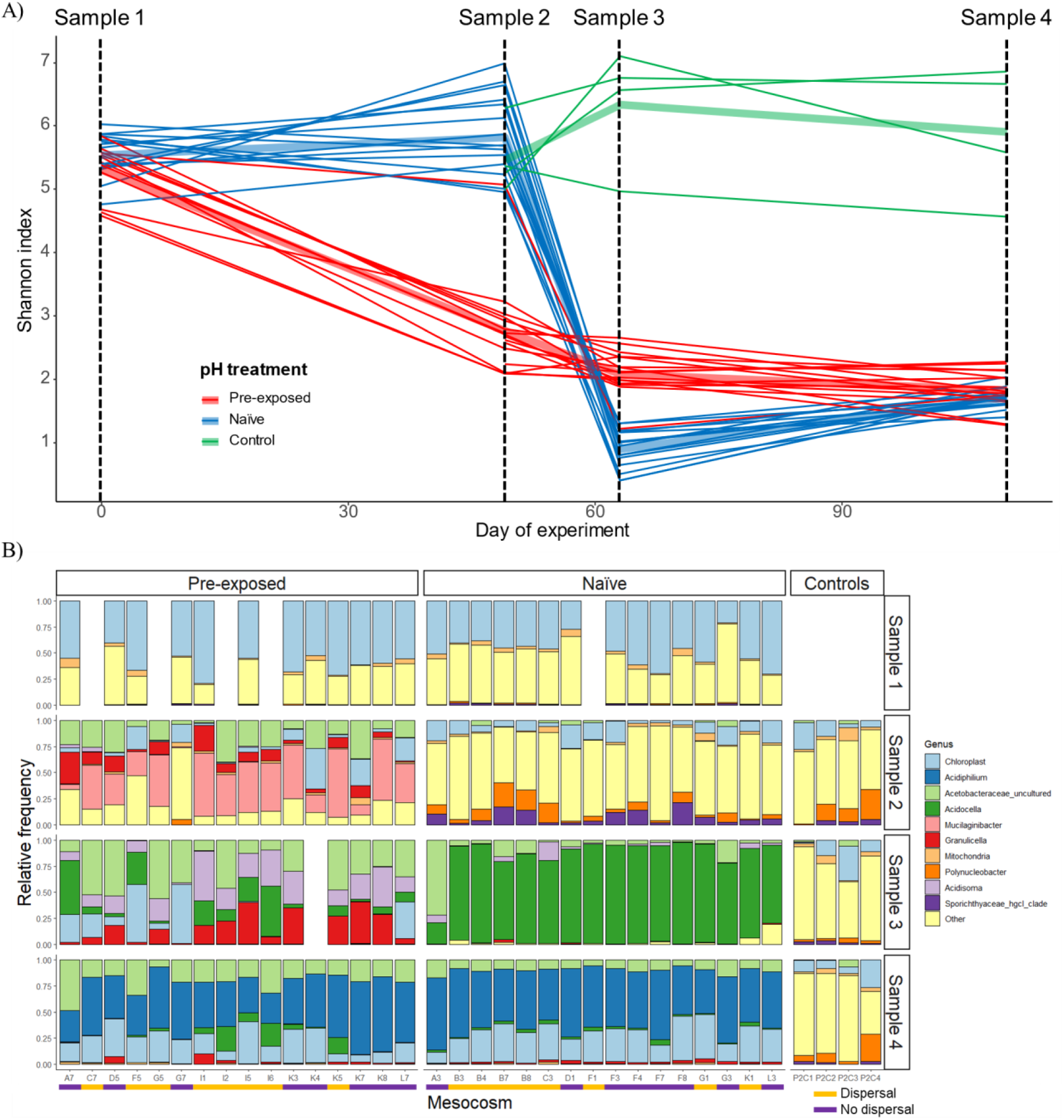
Pre-exposure caused significant changes in alpha diversity and community composition. A) Shannon index values for pre-exposed (red), naïve (blue), and control (green) communities over time. Black dashed lines mark the four time points during the experiment when samples were taken (June 7/day 0, July 26/day 49, August 9/day 63, and September 25/day 111) referred to as Sample 1, Sample 2, Sample 3, and Sample 4. B) Genus-level taxonomic composition of pre-exposed, naïve, and control communities. The top ten genera are colored individually, and all others are grouped together in yellow. Phase I dispersal treatment indicated by orange (dispersal) and purple (no dispersal) lines.

### Distinct ecological dynamics of dominant acidophilic bacteria in pre-exposed versus naïve communities

Beyond changes to alpha diversity, we also investigated how the ecological effects of acidification pre-exposure influenced community composition. Community composition of mesocosms shifted drastically due to acidification in phases I and II (Figure 2B). All communities began with a large diversity of bacteria, primarily from the Bacteroidota and Proteobacteria phyla (representing 93.3% of bacterial ASVs at the start of the experiment). Pre-exposed communities became dominated by the family Acetobacteraceae and genera *Mucilaginibacter* and *Granulicella* by the end of phase I (Sample 2). In contrast, naïve communities remained highly diverse, with an increase in the genus *Polynucleobacter* and the family Sporichthyaceae, which were also present at relatively high frequencies in control communities. Dispersal did not affect species composition in either pre-exposed or naïve communities (Figure 2B). Immediately after severe acidification in phase II (Sample 3), communities within pre-exposed mesocosms continued to shift, with an increase in *Acidocella* and *Acidosoma* and the disappearance of most minor genera. In sharp contrast, composition of naïve communities collapsed to a single genus, *Acidocella*, which overwhelmingly dominated (Figure 2B). Although pre-exposed and naïve communities were distinct in the initial period after severe acidification, by the end of the experiment (Sample 4), regardless of pre-exposure and dispersal, all communities converged and became dominated by *Acidiphilium*, followed by Acetobacteraceae, *Granulicella*, and *Acidocella*. In comparison, control communities not acidified in phase II showed no clear change in community composition over time (Figure 2B).

### Divergence followed by convergence in the evolutionary trajectories of *Acidiphilium rubrum* in pre-exposed and naïve communities

Next, we tested the hypothesis that following severe acidification, the evolutionary trajectories of the dominant species, *Acidiphilium rubrum*, would differ depending on whether it had experienced pre-exposure. Metagenomic shotgun sequencing produced a final contig database of ∼3.1 million contigs totaling approximately 20.8 Gb, which were binned into 81 metagenome-assembled genomes (MAGs) that included *A. rubrum* and *Acidocella* also present in the 16S rRNA gene sequencing data (Supplementary Text). To characterize the impact of pre-exposure on the genetic diversity within *A. rubrum*, we mapped reads to the *A. rubrum* reference genome (NCBI RefSeq assembly GCF_900156265.1, strain ATCC 35905) to call single nucleotide polymorphisms (SNPs) for population genetic analyses. At the start of the experiment, *A. rubrum* across communities were genetically similar, but by the end of phase I (Sample 2), they had diverged significantly both from starting populations as well as between pre-exposed and naïve communities, as quantified by the *F_ST_* metric of population differentiation (Figure 3). We also observed differentiation among certain naïve communities by the end of phase I (Sample 2) (Figure S4). By the end of the experiment (Sample 4), *F_ST_* of *A. rubrum* between almost all pre-exposed and naïve communities decreased, indicating that population genetic diversity had converged after initial divergence (Figure 3, Figure S4). Pairwise *F_ST_* of *A. rubrum* among communities over time ranged between 0 and 1 (Figure S4).

**Figure 3.**
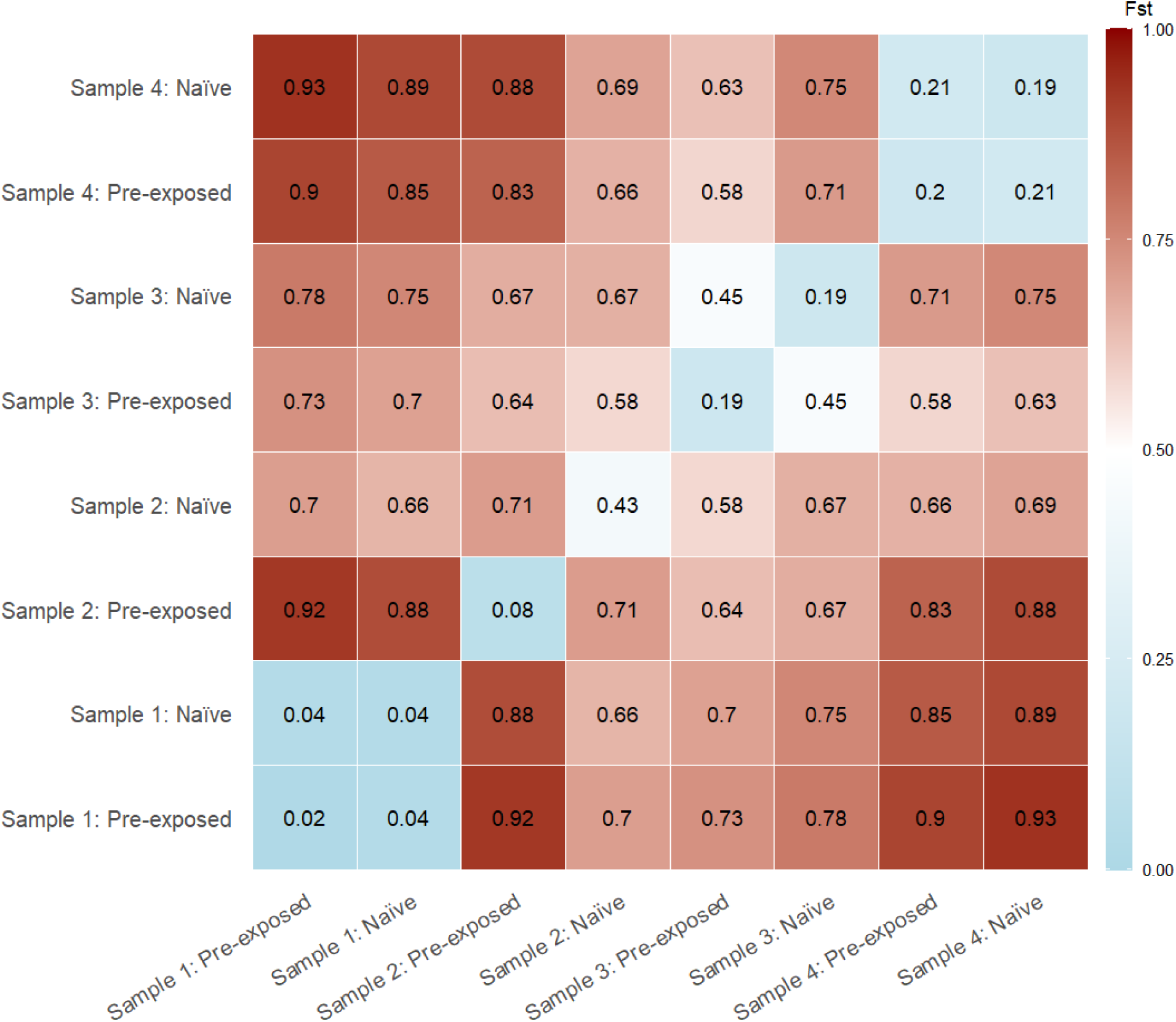
Average pairwise fixation index (*F_ST_*) of SNPs within *Acidiphilium rubrum*. *F_ST_* values are listed chronologically from left to right and from bottom to top (Sample 1 – Sample 4) where values range from 0 (light blue) to 1 (dark red).

### Parallel evolution and the maintenance of genetic diversity in pre-exposed acidophilic bacteria

To assess changes in the genetic diversity of the species remaining at the end of the experiment, we mapped reads to a custom genome database created by combining assembled MAGs with reference genomes of *Acidiphilium*, *Acidocella*, and *Granulicella* species (Supplementary Text). We measured how major alleles in these species shifted over time by calculating genome-wide polarized major allele frequency change (subtracting the frequency of each major allele by the frequency of that same allele at the next time point). Despite large heterogeneity in read mapping across different species, changes in SNP allele frequencies showed similar patterns within *Acidiphilium* and *Acidocella* (Figure 4). For pre-exposed communities, all five *Acidiphilium* and all three *Acidocella* species exhibited significantly greater change in allele frequencies one week after severe acidification (Sample 2 – Sample 3) than the subsequent eight weeks of sustained severe acidification (Sample 3 – Sample 4). Additionally, except for *A. iwatense* and *A.* sp. *KAb 2-4*, species in pre-exposed communities exhibited less change in allele frequencies than naïve communities in those eight weeks (Sample 3 – Sample 4).

**Figure 4.**
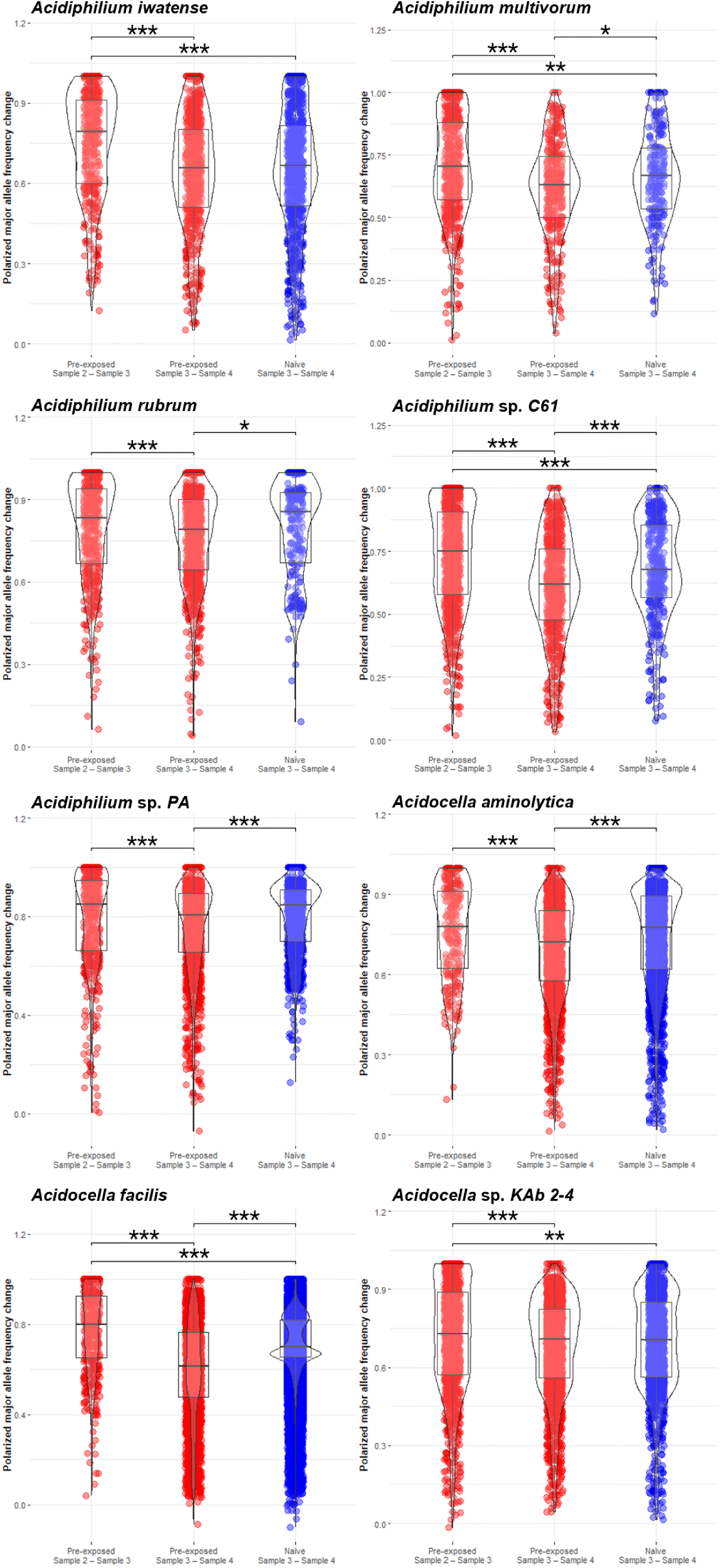
Allele frequency change of single nucleotide polymorphisms (SNPs) polarized to major alleles across species. We calculated allele frequency change by subtracting the frequency of the major allele by the frequency of that same allele in a subsequent time point. Colors indicate pre-exposed (red) and naïve (blue) communities. Asterisks indicate statistical significance based on Holm-Bonferroni adjusted *P*-values from Dunn’s test (*: *P* < 0.05, **: *P* < 0.01, ***: *P* < 0.001).

Because *A. rubrum* was by far the most dominant species at the end of the experiment (Figures S5–S6), we further assessed its evolutionary trajectory and parallelism. We hypothesized that pre-exposed *A. rubrum* populations would maintain greater genetic diversity during severe acidification, as reflected by lower average nucleotide identity (ANI) and higher nucleotide diversity (π). In support of this hypothesis, mean pairwise ANI of *A. rubrum* was significantly lower in pre-exposed than naïve communities at both time points after severe acidification (Sample 3: Z = 2.93, *P* < 0.05; Sample 4: Z = 2.82, *P* < 0.01, Holm-Bonferroni adjusted *P*-values from Dunn’s test) (Figure 5A) and π was also significantly higher in pre-exposed than naïve communities (Sample 3: Z = −4.24, *P* < 0.001; Sample 4: Z = −3.19, *P* < 0.001) (Figure 5B). Additionally, we found evidence of parallel genetic adaptation to acidification among *A. rubrum* populations. Of all SNPs present after severe acidification (present at both Sample 3 and Sample 4), 36 SNPs were shared among pre-exposed communities, 31 SNPs were shared among naïve communities, and only 12 SNPs were shared across populations in pre-exposed and naïve communities (Figure 6A). Permutation tests indicated that the number of shared SNPs among the pre-exposed and naïve communities were significantly higher than neutral expectations when SNPs were randomized across mesocosms (pre-exposed: *P* < 0.05; naïve: *P* < 0.05, N = 10,000) (Figure 6B).

**Figure 5.**
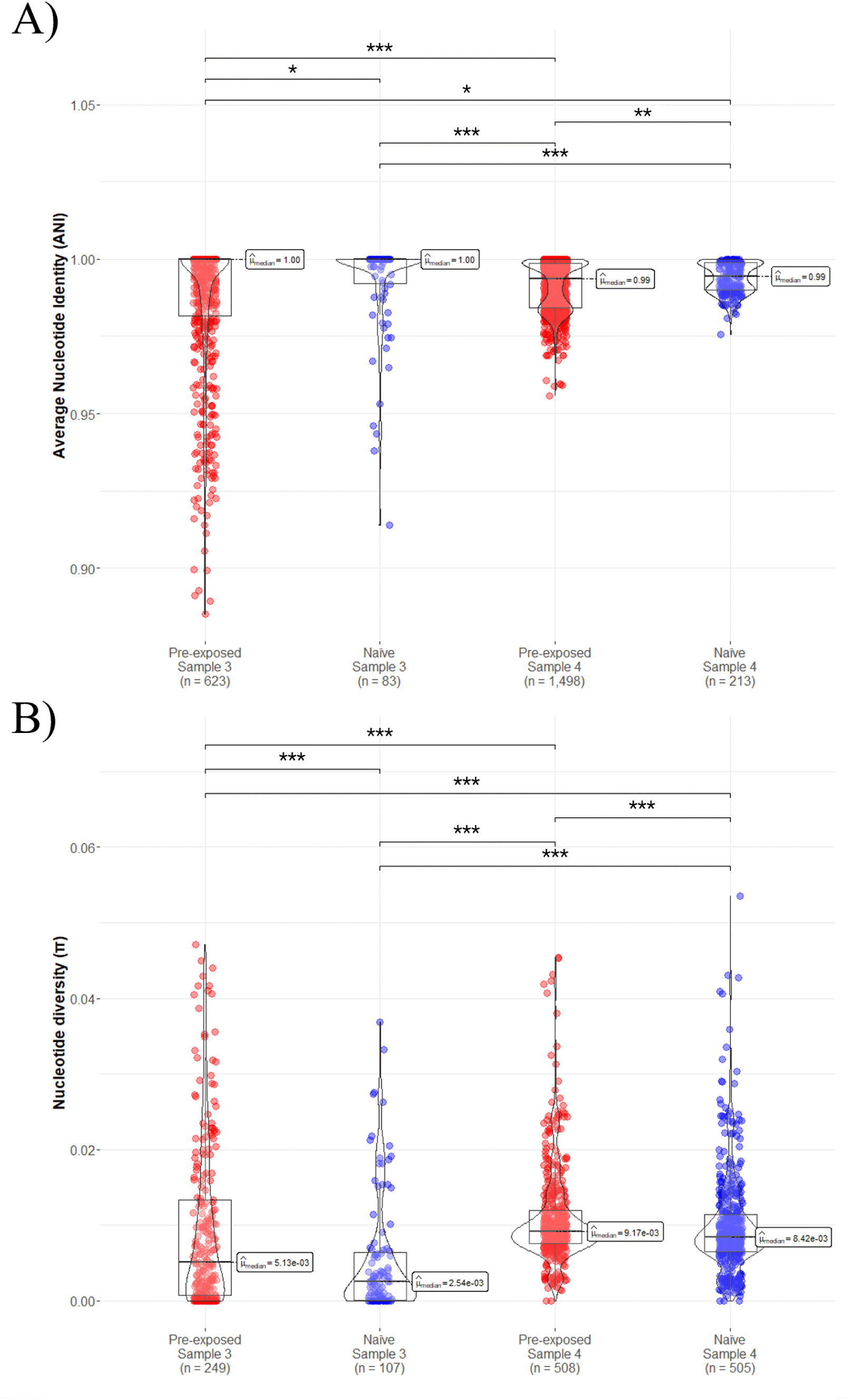
Effects of pre-exposure on evolution of *Acidiphilium rubrum*. A) Pairwise average nucleotide identity (ANI) and B) nucleotide diversity (π) of *A. rubrum* genomes within pre-exposed (red) and naïve (blue) communities after one week (Sample 3) and eight weeks (Sample 4) of severe acidification. Number of scaffolds are in parentheses. Asterisks indicate statistical significance based on Holm-Bonferroni adjusted *P*-values from Dunn’s test (*: *P* < 0.05, **: *P* < 0.01, ***: *P* < 0.001).

**Figure 6.**
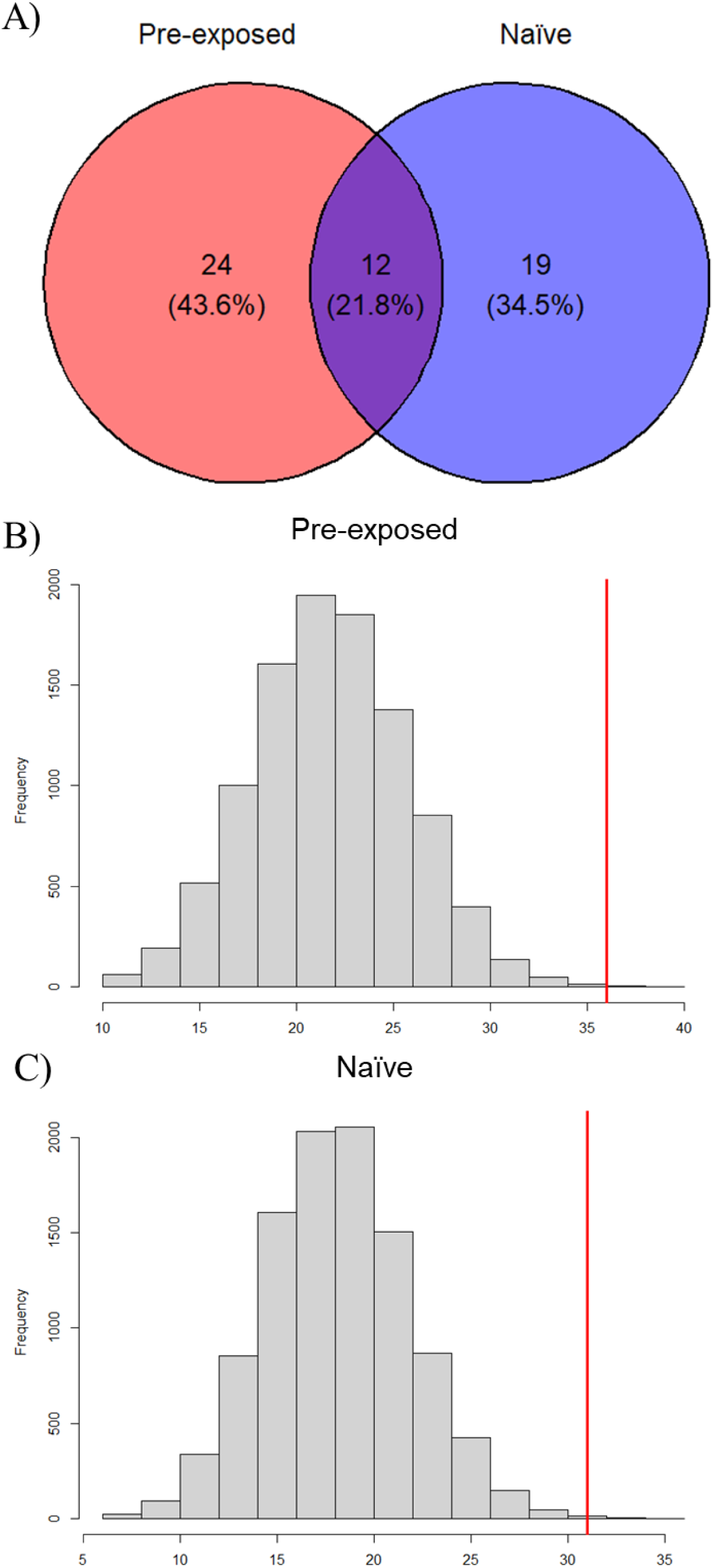
Shared single nucleotide polymorphisms (SNPs) between and among *Acidiphilium rubrum* populations. A) The number of shared SNPs between and among pre-exposed and naïve communities consistently present after severe acidification (at both Sample 3 and Sample 4). Permutations of mesocosms over SNPs indicate that the number of shared SNPs within B) pre-exposed and C) naïve communities are significantly greater than neutral expectations (pre-exposed: *P* < 0.05; naïve: *P* < 0.05, N = 10,000). Red lines indicate the observed number of shared SNPs.

In summary, two major evolutionary patterns emerged: 1) in pre-exposed populations, the greatest genetic change occurs immediately after severe acidification, 2) following long-term severe acidification, genetic change was significantly less in pre-exposed than in naïve populations. These patterns are consistent with pre-exposure to moderate acid stress causing parallel directional selection for acid-tolerant alleles, resulting in more resistant communities that experience less dramatic population bottlenecks when confronted with severe acidification, thereby retaining greater genetic diversity.

## Discussion

Environmental stress can negatively impact community diversity and ecosystem functioning, but pre-exposure to sublethal levels of stress could mediate their persistence and recovery. Investigating the contributions of species sorting and evolutionary adaptation to this process deepens our understanding of the underlying mechanisms of community recovery during sustained stress. Here, we provide evidence that pre-exposure to moderate acidification altered the response of aquatic microbial communities during severe acidification, both ecologically through changes in community composition and evolutionarily via genome-wide shifts of genetic variation across multiple species.

### Species sorting reshapes community diversity under stress

Pre-exposure to acidification had profound effects on bacterial communities through species sorting as indicated by significantly lower Shannon index and markedly different relative abundances of genera. Although pre-exposure was initially detrimental to alpha diversity, pre-exposed communities exhibited relatively greater resistance to severe acidification; Shannon diversity not only decreased significantly less than naïve communities but also exhibited significantly higher absolute values immediately after severe acidification (Sample 3).

This initial relative increase of community resistance in pre-exposed communities against the immediate effects of severe acidification (between Sample 2 and Sample 3) may be due in part to species sorting that favored acidophiles previously at very low frequencies at the start of the experiment, such as the family Acetobacteraceae, which contains several acidophilic genera including *Acidiphillium*, *Acidisoma*, and *Acidocella* ^39^. Pre-exposure also selected for *Granulicella*, a genus of acidophiles within the family Acidobacteriaceae ^40^, and *Mucilaginibacter*, a diverse genus within the family Sphingobacteriaceae that contains species previously isolated from acidic forest soils ^41^ and documented to grow in acidic conditions as low as pH 2 ^42^. In contrast, these taxonomic groups were not detected in naïve communities at the end of phase I (Sample 2) due to either low abundance or absence. Instead, there was an increase in the genus *Polynucleobacter*, a ubiquitous and diverse freshwater bacterioplankton that can tolerate a wide range of environmental conditions ^43–46^, and the family Sporichthyaceae, which contains four named species of motile facultative anaerobes with aerial hyphae isolated from soil, lake sediment, and human skin ^47–51^. Both *Polynucleobacter* and Sporichthyaceae were also observed in control communities at the end of pre-exposure (Sample 2), so their presence likely indicates seasonal effects or selection by the mesocosm environment.

Immediately after severe acidification, alpha diversity within naïve communities crashed and became overwhelmingly dominated by *Acidocella* with minor contributions from Acetobacteraceae, *Acidisoma*, and *Granulicella* in some but not all mesocosms. Notably, Acetobacteraceae and *Granulicella* were among the selected taxa in pre-exposed communities at the end of pre-exposure (Sample 2). Contrary to our hypothesis, pre-exposure did not improve long-term community resistance, with both Shannon indices and community composition ultimately converging across pre-exposed and naïve communities by the end of the experiment, perhaps because responses of bacterial communities to stress are rapid enough to occur over a few weeks even in communities with no history of exposure to a stressor. However, despite this convergence, pre-exposure did cause a significantly different species sorting trajectory immediately after severe acidification (Sample 3), thus providing supporting evidence that pre-exposure to a weaker dose of a stressor could allow more time for both plastic and evolutionary responses by resistant organisms.

Shannon diversity consistently decreased in pre-exposed mesocosms throughout the experiment, albeit at lower rates between each successive time point. In contrast, alpha diversity and richness (observed number of ASVs) in naïve communities did recover significantly under severe acidification (Sample 3 – Sample 4) to eventually reach the same levels as pre-exposed communities by the end of the experiment. Although the 16S rRNA analysis included mesocosms under dispersal, we did not observe any significant effects of dispersal on alpha diversity or richness at any time point. Furthermore, there was no additional input from the source lake to mesocosms after the start of the experiment so the increase in the number of observed ASVs during recovery is unlikely to be attributable to migration. Thus, seemingly novel ASVs at the end of the experiment such as those assigned to the genus *Acidiphilium* were most likely previously present, but at undetectably low absolute or relative abundances, and would have required sufficient positive selection to overcome loss through ecological drift. Our findings are consistent with a previous study on the same source lake that successfully recovered non-obligate acidophilic bacteria capable of surviving at pH 2, which suggests that such acidophiles are ever present in ordinary freshwater, even from protected water sources ^52^. Recovery of alpha diversity in naïve communities under severe acidification (Sample 3 – Sample 4) was therefore driven by a combination of population growth of initially undetected acidophiles and increases in taxonomic evenness.

All communities besides controls converged to a single profile composed of mostly *Acidiphilium* at the end of the experiment. *Acidiphilium* (meaning “acid lover”) is a genus of gram negative, motile, flagellated, photosynthetic, straight rod Proteobacteria containing eight named species ^53–55^. *Acidiphilium* are known to be mesophilic and obligately acidophilic, growing between pH 2.0–5.9 but not above 6.1 ^55^. Here, we show that while *Acidiphilium* may not grow well under neutral pH conditions, it does persist at low levels in natural lake freshwater of approximately pH 8.5 and can rapidly increase if environmental conditions become sufficiently acidified. Putative acidophiles selected through species sorting in pre-exposed communities (Acetobacteraceae and *Granulicella*) also persisted at the end of the experiment and at slightly greater frequencies than in naïve communities, supporting the hypothesis that pre-exposure has long-term effects on species composition.

### Evolutionary responses to acidification

We identified and tracked SNPs across time in nine reference genomes (five *Acidiphilium* species, three *Acidocella* species, and *Granulicella* sp. *5B5*) and a single MAG annotated as an unknown Rickettsiales bacterium. Significant changes in genome-wide allele frequencies, including SNPs where the major allele was completely replaced, were observed in all genomes between at least two time points, demonstrating the role of evolutionary processes in community responses to acidification. Although we cannot definitively conclude that adaptation played a causal role in community responses (*i.e.*, evolutionary and ecological responses could be occurring at least partially independently), we did observe significant evolutionary change within successful acidophiles, which may have contributed to their persistence. Because these genomes were not detected in control communities, we were unable to characterize and directly compare SNPs from populations that did not experience severe acidification. However, of the six other MAGs that mapped reads from control communities, genome-wide allele frequency changes were significantly lower, as expected in the absence of a stressor (Figure S7).

We tracked SNPs in pre-exposed communities at the onset of severe acidification (Sample 2 – Sample 3) as well as in both pre-exposed and naïve communities throughout the eight weeks of severe acidification (Sample 3 – Sample 4) across all detected species of *Acidiphilium* and *Acidocella*. These eight acidophilic species independently exhibited strikingly similar patterns of genome-wide allele frequency changes (Figure 4). In pre-exposed communities, significantly greater change was observed one week after severe acidification (Sample 2 – Sample 3) than the following eight weeks until the end of the experiment (Sample 3 – Sample 4). In those eight weeks of severe acidification, six of the eight acidophilic species (excluding *A. iwatense* and *A.* sp. *KAb 2-4*) exhibited significantly lower change in genome-wide allele frequencies in pre-exposed communities than naïve communities. Importantly, this shows that the magnitude of evolutionary response is largest upon initial exposure despite the stressor being applied at a lower level. This is consistent with these evolutionary changes representing the beginning of an ‘adaptive walk’ towards a new fitness optimum, when theory predicts changes will be the greatest ^56^. Thus, pre-exposure served to initiate adaptation earlier, with the response diminished upon subsequent exposure to greater levels of stress. This may be viewed as analogous to the response to vaccination, where exposure to a weak dose of virus permits an immune response that reduces the impact of exposure to greater viral loads in the future.

### The dominance of Acidiphilium rubrum

16S rRNA and MAG analysis indicated that by the end of the experiment, all communities, regardless of pre-exposure, were dominated by a single genus, *Acidiphilium* (with minimal contributions by *Acidocella*). Competitive metagenomic read mapping of *Acidiphilium* reference genomes revealed that *Acidiphilium rubrum* was largely responsible for this dominance, followed by *Acidiphilium* sp. *PA* (Figure S6). *A. rubrum* is a highly acidophilic purple bacterium that can be isolated from acid mine drainage sites of pH 2–3 ^57–59^. The ascendancy of *A. rubrum* suggests that this species has a selective advantage over other acidophiles at such low pH. We observed significant differentiation between *A. rubrum* in pre-exposed and naïve communities at the end of pre-exposure (Sample 2: *F_ST_* = 0.71), followed by convergence by the end of the experiment (Sample 3: *F_ST_* = 0.45, Sample 4: *F_ST_* = 0.21), indicating independent evolutionary trajectories that eventually reached the same destination (Figure 3). Interestingly, despite the absence of experimental manipulation in phase I, *A. rubrum* populations in naïve communities exhibited substantial genetic divergence (Sample 1 – Sample 2: *F_ST_* = 0.66), albeit significantly less than in pre-exposed populations (Sample 1 – Sample 2: *F_ST_* = 0.92).

Further population genetic analysis of *A. rubrum* populations provides additional evidence for the effects of pre-exposure on evolutionary processes. Pairwise ANI and π reveal that pre-exposure allowed *A. rubrum* to retain significantly greater genetic diversity at both time points after severe acidification, potentially because pre-exposure induced directional selection, producing more acid-tolerant populations that underwent less severe bottlenecks when confronted by severe acidification (Figure 5). By the end of the experiment, genetic diversity of *A. rubrum* increased significantly relative to the onset of severe acidification in all communities, but the effect of pre-exposure was still evident as *A. rubrum* populations in pre-exposed communities were significantly more diverse than naïve communities. The impact of pre-exposure on *A. rubrum* evolution can also be observed through the number of individual SNPs shared exclusively among either pre-exposed or naïve populations. Significantly more SNPs were shared among *A. rubrum* populations within pre-exposed and within naïve communities than under neutral expectations, which suggests that pre-exposure independently selected for alleles not under selection in naïve communities (Figure 6). In summary, the high number of parallel SNP changes suggests that severe acidification caused strong selection, and the distribution of shared SNPs suggests that distinct SNPs were targeted in pre-exposed and naïve communities. While it may be counter-intuitive that selection due to pre-exposure can prevent loss of genetic diversity, our study indicates that pre-exposed communities may suffer less dramatic population bottlenecks when confronted with stronger selection, which can ultimately help maintain genetic diversity.

### Conclusions

In this study, we show that pre-exposure to environmental stress can help biological communities maintain taxonomic and genetic diversity against future severe stress. Despite the convergence of community composition between pre-exposed and naïve communities, and despite the loss of many taxa in all acidified communities, we demonstrate that species sorting due to pre-exposure generated greater community resistance and mitigated declines in taxonomic diversity. Additionally, we show that pre-exposure caused independent evolutionary processes that resulted in reduced genome-wide change following severe acidification and greater levels of genetic diversity in the species that dominated acidified environments. Thus, we provide evidence for the dual roles of species sorting and evolutionary adaptation in community responses to severe stress, as well as the importance of pre-exposure for adaptation ^10,12,60–63^.

## Methods

### Study site

We conducted the experiment at the Large Experimental Array of Ponds (LEAP) facility located at the Gault Nature Reserve in Mont-Saint-Hilaire, Quebec, Canada. We filled replicate 1,000 L mesocosms on May 23/24, 2017, by sourcing water from the nearby Lake Hertel (45°32’ N, 73°09’ W), which is protected under UNESCO as part of the Mont Saint Hilaire Biosphere Reserve. The naturally mesotrophic lake has a maximum depth of 8.2 m and a natural pH of 7.5– 9.5 ^64,65^. We used a coarse sieve to filter water from Lake Hertel before it entered the mesocosms, which prevented introduction of fish and large invertebrates leaving a community of naturally co-occurring zooplankton, phytoplankton, bacteria, and viruses.

### Experimental design

We designed the biphasic pulse-press experiment to test the isolated and interacting effects of several levels of acidification pre-exposure and dispersal regimes on community response to severe acidification, which has been described in a previous study ^21^. Here, we focused on only the 16 mesocosms pre-exposed to the strongest acidification treatment of pH 4 as well as the 16 naïve mesocosms that were left untreated and remained at their natural acidity of approximately pH 8.5 (Figure 1A). In phase I of the experiment, we maintained pre-exposure to pH 4 through weekly pulse titration with 10N H_2_SO_4_ for seven weeks, from June 14 – August 2. Pre-exposed mesocosms exhibited a sharp decrease in pH buffering capacity with each acidification pulse in the first weeks of phase I (Figure 1B). Half of pre-exposed and naïve mesocosms were also under a global dispersal regime where we mixed 1% of water from each metacommunity of four mesocosms in a pool and then redistributed it on a weekly basis allowing for migration within metacommunities. We initiated phase II on August 2 when all mesocosms were acidified to pH 3 in a sustained press treatment and dispersal regimes were terminated. Phase II lasted for approximately eight weeks until the end of the experiment on September 25. We also established four isolated control mesocosms subjected to neither phase I nor phase II treatments.

### Sample collection

We monitored mesocosms weekly for water pH (Figure 1B) using multiparameter sondes (Yellow Spring Instruments, Ohio). We used integrated water samplers made from 2.5 cm diameter PVC tubing to sample water biweekly from the top 35 cm of the water column at five random locations within each mesocosm until a total of 2 L of water was collected. We subsequently stored water samples in dark, triple-washed Nalgene bottles at 4°C before filtration later that same day. We used independent samplers and dark bottles for each mesocosm to minimize cross-mesocosm contamination. For each sample, we filtered 500 mL of water at an on-site lab using 0.22 µm pore size, 47 mm diameter Millipore hydrophilic polyethersulfone membranes (Sigma-Aldrich). We then transported filters to the laboratory on dry ice and stored them at −80 °C prior to DNA extraction.

### DNA extractions

We extracted DNA from samples collected across four time points: 1. At the beginning of the experiment one week prior to pre-exposure treatment (June 7/day 0), 2. At the end of phase I after six weeks of pre-exposure (July 26/day 49), 3. At the beginning of phase II one week after severe acidification (August 9/day 63), and 4. At the end of the experiment after approximately eight weeks of severe acidification (September 25/day 111) hereafter referred to as Sample 1, Sample 2, Sample 3, and Sample 4 (Figure 1B). In total, there were 128 samples (16 mesocosms X 2 treatments X 4 time points) and 12 controls (4 control mesocosms X 3 time points excluding Sample 1). We extracted and purified total genomic DNA from half filter papers using the DNeasy PowerWater kit (QIAGEN) following QIAGEN guidelines including a 5-minute vortex of the filter with beads and an additional incubation of 30 minutes at 37°C with 1 μL Rnase (Thermo Scientific) after cell lysis and before the first supernatant transfer to remove RNA contamination ^66^.

### 16S rRNA gene sequencing

We profiled bacterial community composition using 16S rRNA amplicon sequencing. Specifically, we used the primers U515_F (5’-ACACGACGCTCTTCCGATCTYRYRGTGCCAGCMGCCGCGGTAA-3’) and E786_R (5’-CGGCATTCCTGCTGAACCGCTCTTCCGATCTGGACTACHVGGGTWTCTAAT-3’) to target an approximately 200 bp amplicon of the V4 region of the 16S rRNA gene as described previously (Preheim *et al.* 2013). We treated samples that initially failed to PCR amplify with sodium acetate and then ethanol precipitated with GenElute LPA Linear Polyacrylamide (Sigma-Aldrich) to increase DNA concentration ^67^. Genomic DNA quality control, sequencing library preparation, two-step PCR ^66^, and amplicon sequencing via Illumina MiSeq v2 PE250 was conducted at the McGill Genome Centre. An average of 28,387 (range: 0–85,039) 16S rRNA reads were produced per sample. Rarefaction retained 134 (95.71%) samples after removing six samples that produced less than 5 reads.

### Amplicon sequence analysis

We processed raw 16S rRNA amplicon sequences using the QIIME2 bioinformatics pipeline ^68^. We first removed primer sequences using cutadapt followed by identification of ASVs using DADA2 ^69,70^. We aligned ASVs using MAFFT and constructed phylogenetic trees using FastTree 2 based on Jukes-Cantor distances ^71,72^. We created a custom reference database by using the U515_F/E786_R primers to *in silico* extract the target 16S rRNA V4 region from the SILVA 138 database ^73^.

We generated taxonomic weights according to occurrence records in freshwater habitats using redbiom and Qiita by limiting sample type to “fresh water” or “freshwater” and context to “Deblur_2021.09-Illumina-16S-V4-90nt-dd6875” ^74–76^. We used a total of 6,206 V4 16S rRNA sequences (6,003 “fresh water” and 203 “freshwater”) to weight taxonomic assignment towards those found previously in freshwater environments. We then assigned taxonomies to ASVs with a naïve Bayes classifier trained using scikit-learn on the extracted SILVA 138 database that was modulated by the freshwater taxonomic weights ^77^. We accepted taxonomic assignments if classification confidence was at least 0.7 ^78^.

We assessed alpha diversity using the natural logarithm Shannon index computed after we rarified ASVs of each sample to a depth of 1,000 based on saturation of rarefaction curves ^79^. We compared Shannon values between pre-exposed and naïve communities at each time point using Kruskal-Wallis tests ^80^. We also assessed longitudinal differences in Shannon diversity using Wilcoxon signed-rank and Mann-Whitney U tests and statistical significance via Benjamini & Hochberg corrected q-values ^81–84^.

### Metagenomic shotgun sequencing

We selected samples from all isolated (*i.e.,* no dispersal in phase I) pre-exposed and naïve mesocosms except for one mesocosm from each phase I treatment for further metagenomic analysis along with control mesocosms. In total, we subjected 68 samples across the four time points to deep sequencing at an average of 220 million reads per sample. We focused sequencing on phase I samples (Sample 1 and Sample 2 at ∼330 million reads/sample) compared to phase II samples (Sample 3 and Sample 4 at ∼110 million reads/sample) to maximize the probability of detecting and quantifying genetic diversity within dominant phase II species that were initially at low abundances in phase I. Quality control, library prep, and sequencing on Illumina NovaSeq 6000 PE150 were conducted at the McGill Genome Centre.

### Metagenomic analysis

We processed and analyzed metagenomic sequences within the anvi’o framework ^85^. We first removed Illumina TruSeq LT adaptors with cutadapt and quality filtered reads using illumina-utils ^86^. We used MEGAHIT to co-assemble reads from the same mesocosm across the four time points ^87^. We merged all contigs and removed those less than 2,500 bp. We then mapped reads from each sample to contigs using Bowtie2 and SAMtools ^88,89^. We identified prokaryotic genes in the contigs using Prodigal ^90^. We used hidden Markov models for collections of 71 bacteria, 76 archaea, and 83 protist single-copy core genes (SCGs) to identify and recover them from contigs ^91–93^.

We clustered contigs into bins using CONCOCT and MetaBAT2, which we then dereplicated and aggregated into metagenome-assembled genomes (MAGs) using DAS Tool ^94–96^. We estimated completeness and redundancy of MAGs based on SCG collections. We determined prokaryotic taxonomic identities of MAGs via the most frequent of 22 bacterial SCGs from the Genome Taxonomy Database (GTDB) using DIAMOND ^97,98^. We included MAGs with a completeness >90% and redundancy <10% for further analysis along with the most dominant MAG at the end of the experiment (Sample 4) based on mean coverage within samples from each phase I treatment and time point. For the dominant MAG at the end, we used a reference genome approach to identify single nucleotide polymorphisms (SNPs) occurring in at least two samples and calculated pairwise fixation index (*F_ST_*) within anvi’o ^99,100^.

Three genera (*Acidiphilium*, *Acidocella*, and *Granulicella*) dominated phase II (Sample 3 and Sample 4) communities based on 16S rRNA gene amplicon and MAG results. We further assessed population microdiversity of species within these genera using InStrain ^101^. We downloaded reference genomes from the three genera from NCBI RefSeq and merged them with MAGs to create a custom genome database that we then dereplicated using dRep, checkM, MASH, and fastANI with a MASH sketch size of 10,000 and a minimum overlap between genomes of 0.3 ^102–106^. We used Prodigal to profile genes for each genome, and we competitively mapped reads against reference genomes and MAGs using Bowtie2 and SAMtools ^88–90^. We called SNPs using a minimum coverage threshold of 5 and a minimum SNP frequency of 0.05 using InStrain ^101^. We also used InStrain to calculate scaffold-level metrics including ANI and π ^101,107,108^. We calculated allele frequency change of polarized major alleles through subtracting the frequency of the major allele at each SNP by the frequency of that same allele in a subsequent time point. We longitudinally compared pairwise ANI and π as well as allele frequency changes between pre-exposed and naïve communities using Dunn’s test and assessed statistical significance via Holm-Bonferroni adjusted p-values ^109,110^. For the most dominant species identified at the end of the experiment (*A. rubrum*), we used permutation tests randomizing the mesocosm of each SNP to assess the significance of the number of shared SNPs between populations in pre-exposed and naïve communities. We only considered SNPs consistently present after severe acidification (at both Sample 3 and Sample 4) as a conservative approximation for adaptation.

## Acknowledgements

We are grateful to Charles Bazerghi, Kristina Krebs, Ilke Geladi, and Natalie Chehab for their efforts in establishing and maintaining the mesocosms during the experiment at the LEAP facility. We would also like to thank Lauren Bennett, Michael Maddalena, Nicole Stinson, Rachel Takasaki, and Kiran Yendamuri for assistance in the lab. High Performance Computing resources were provided by the Digital Research Alliance of Canada through a Resource Allocation Competition grant awarded to R.D.H.B. We are very thankful for the technical support team at the Alliance, especially Huizhong Lu, Jose Sergio Hleap, Pier-Luc St-Onge, Daniel Stubbs, and Denise Koch for their expertise and patience. C.C.Y.X. was funded in part by a Vanier Graduate Student Scholarship. The LEAP facility was built with support of the Canada Foundation for Innovation and the Liber Ero Chair in Biodiversity Conservation. The research was supported by a Discovery Grant from the Natural Sciences and Engineering Research Council of Canada (R.D.H.B., A.G.), the Liber Ero Chair in Biodiversity Conservation (A.G.), the Fond Quebecois de la Recherche – Nature et Technologies, the Canada Research Chairs programme (M.E.C., A.G., B.J.S., R.D.H.B.), the Quebec Centre for Biodiversity Science, and the Groupe de Recherche Interuniversitaire en Limnologie.

## Supplementary Text

### DNA sequencing and mapping

We sequenced an average of 242 million raw read pairs per sample (range: 100–551 million). To maximize probability of detecting very rare acidophiles that became dominant after severe acidification, we sequenced samples from phase I at greater depth compared to samples from phase II (Mean read pairs in millions; Sample 1: 260; Sample 2: 370; Sample 3: 204; Sample 4: 143). On average, 93.77% of read pairs passed quality filtration (227 million read pairs per sample). Each mesocosm co-assembly contained an average of 0.7 million contigs with an average total base count of approximately 1.95 Gb (2,875 bp per contig). An average of 89.58% of read pairs mapped successfully to contigs resulting in an average of 203 million read pairs mapped per sample. A lower proportion of read pairs mapped in samples from the start of the experiment than at any other time point (Sample 1: 75.87%, Sample 2: 89.21%, Sample 3: 91.49%, Sample 4: 93.86%), consistent with a high proportion of reads originating from unassembled eukaryotic algae. The final contig database consisted of about 3.1 million contigs totaling approximately 20.8 Gb, and 2.1 million annotated genes from an estimated 2,328 bacterial and 64 eukaryotic genomes.

### Metagenome-assembled genome (MAG) detection

We binned contigs into 81 MAGs accounting for approximately 400 Mb, which represented 1.90% of all nucleotides. Bacterial MAGs ranged from 524 Kb to 15.6 Mb and varied in quality (52.11–100% completeness, 0–85.92% redundancy). Two eukaryotic MAGs were also identified (23.95 Mb and 19.14 Mb). Twenty-two MAGs with a minimum of five bacterial single-copy core genes (SCGs) were taxonomically classified to the phyla Actinobacteriota, Bacteroidota, Proteobacteria, and Verrucomicrobiota based on at least 75% supporting SCGs (Table S2). These corresponded to 14 unique genera and 12 species, including *Acidiphilium rubrum* and *Acidocella*, which were also present in the 16S rRNA gene sequencing data. The MAG assigned to *A. rubrum* constituted 97–98% of all mapped reads at the end of the experiment (Sample 4) across both pre-exposed and naïve communities.

### Mapping to custom reference database

We obtained 44 reference genomes from 17 species within *Acidiphilium*, *Acidocella*, and *Granulicella* from NCBI RefSeq and merged them with MAGs to create a custom genome database, which we dereplicated to 29 reference genomes and 29 MAGs. Reads from pre-exposed and naïve communities mapped to five *Acidiphilium* species (*A. iwatense*, *A. multivorum*, *A. rubrum*, *A.* sp. *C61*, *A.* sp. *PA*), three *Acidocella* species (*A. aminolytica*, *A. facillis*, *A.* sp. *KAb 2-4*), *Granulicella* sp. *5B5*, and a single MAG assigned to an unnamed species within the order Rickettsiales (family SXRF01, genus *RFOF01*). Reads from control communities mapped only to other MAGs.

Mean coverage of genomes differed significantly across species and between pre-exposed and naïve communities across time (Figure S5). Despite higher sequencing depth of samples prior to severe acidification, the total number of mapped reads was dwarfed by samples from the start of phase II (Sample 3) and the end of the experiment (Sample 4) (Figure S6A). Samples from naïve mesocosms at the beginning of the experiment (Sample 1) and at the end of phase I (Sample 2) yielded relatively few mapped reads, mapping only to *A.* sp. *KAb 2-4* (average of 17,875 and 44,997 mapped read pairs per sample resulting in approximately 1X and 2.3X coverage respectively). Reads from pre-exposed communities at the end of phase I (Sample 2) mapped across all 10 species (Figure S6B). Reads from communities immediately after severe acidification (Sample 3) mapped mostly to *A. facilis* with many more mapped from naïve than pre-exposed communities. At the end of the experiment (Sample 4), most reads mapped to *A. rubrum* followed by *A.* sp. *PA*.

---

**Table S1.** Significance and direction of pairwise change in Shannon index values. Bonferroni *P_FDR_*-values based on *P*-values from Wilcoxon signed-rank tests.

|  |  | **W** | **P** | **$P_{FDR}$** | **Direction** |
| --- | --- | --- | --- | --- | --- |
| Pre-exposed | Sample 1 – Sample 2 | 0.0 | 0.000488 | 0.000977 | Negative |
|  | Sample 2 – Sample 3 | 6.0 | 0.000854 | 0.00128 | Negative |
|  | Sample 3 – Sample 4 | 20.0 | 0.0215 | 0.0323 | Negative |
| Naïve | Sample 1 – Sample 2 | 35.0 | 0.169 | 0.169 |  |
|  | Sample 2 – Sample 3 | 0.0 | 0.000031 | 0.000092 | Negative |
|  | Sample 3 – Sample 4 | 0.0 | 0.000031 | 0.000092 | Positive |
| Control | Sample 2 – Sample 3 | 2.0 | 0.375 | 0.375 |  |
|  | Sample 3 – Sample 4 | 3.0 | 0.625 | 0.625 |  |

**Table S2.** Significance of pre-exposure on pairwise change in Shannon index values. Bonferroni *P_FDR_*-values based on *P*-values from Mann-Whitney U tests.

|  | **U** | ***P*** | ***P<sub>FDR</sub>*** |
| --- | --- | --- | --- |
| Sample 1 – Sample 2 | 2.0 | 0.00002 | 0.00002 |
| Sample 2 – Sample 3 | 240.0 | 0.000002 | 0.000007 |
| Sample 3 – Sample 4 | 2.0 | 0.000003 | 0.000010 |

**Table S3.** Significance of dispersal on Shannon index values. Benjamini-Hochberg *Q*-values based on *P*-values from Kruskal-Wallis tests.

|  | **H'** | ***P*** | ***Q*** |
| --- | --- | --- | --- |
| Sample 1 | 0.0219 | 0.882 | 0.896 |
| Sample 2 | 1.195 | 0.274 | 0.351 |
| Sample 3 | 0.400 | 0.527 | 0.604 |
| Sample 4 | 4.455 | 0.0348 | 0.0638 |

**Table S4.**
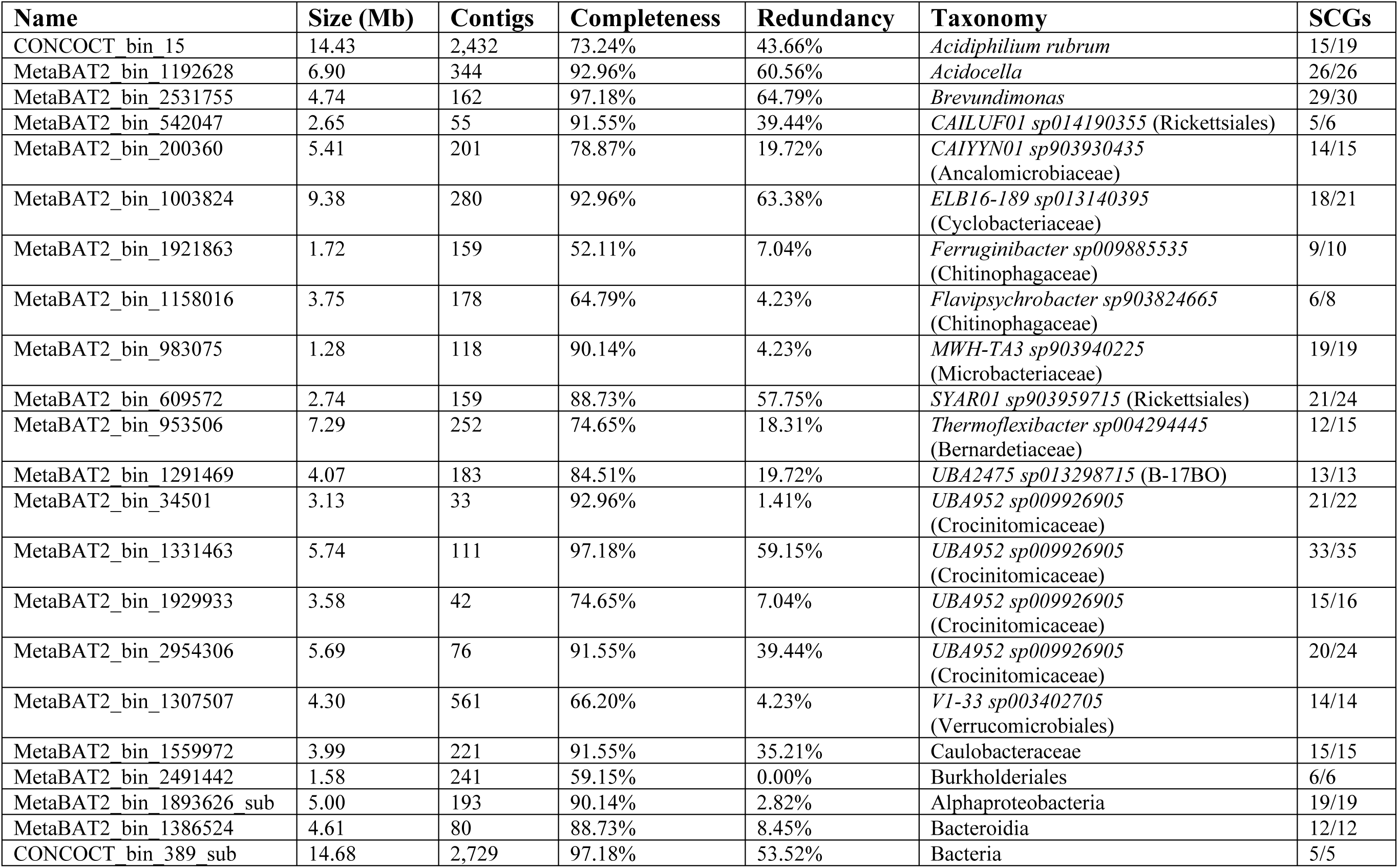
Summary of taxonomically classified metagenome-assembled genomes (MAGs) Taxonomic classifications based on 75% concordance of five minimum single-copy core genes (SCGs) out of 22 unique bacterial SCGs (some MAGs contain more than 22 total SCGs due to redundancy). Genus and species identities shown when available with family/order identity in parentheses.

**Figure S1.**
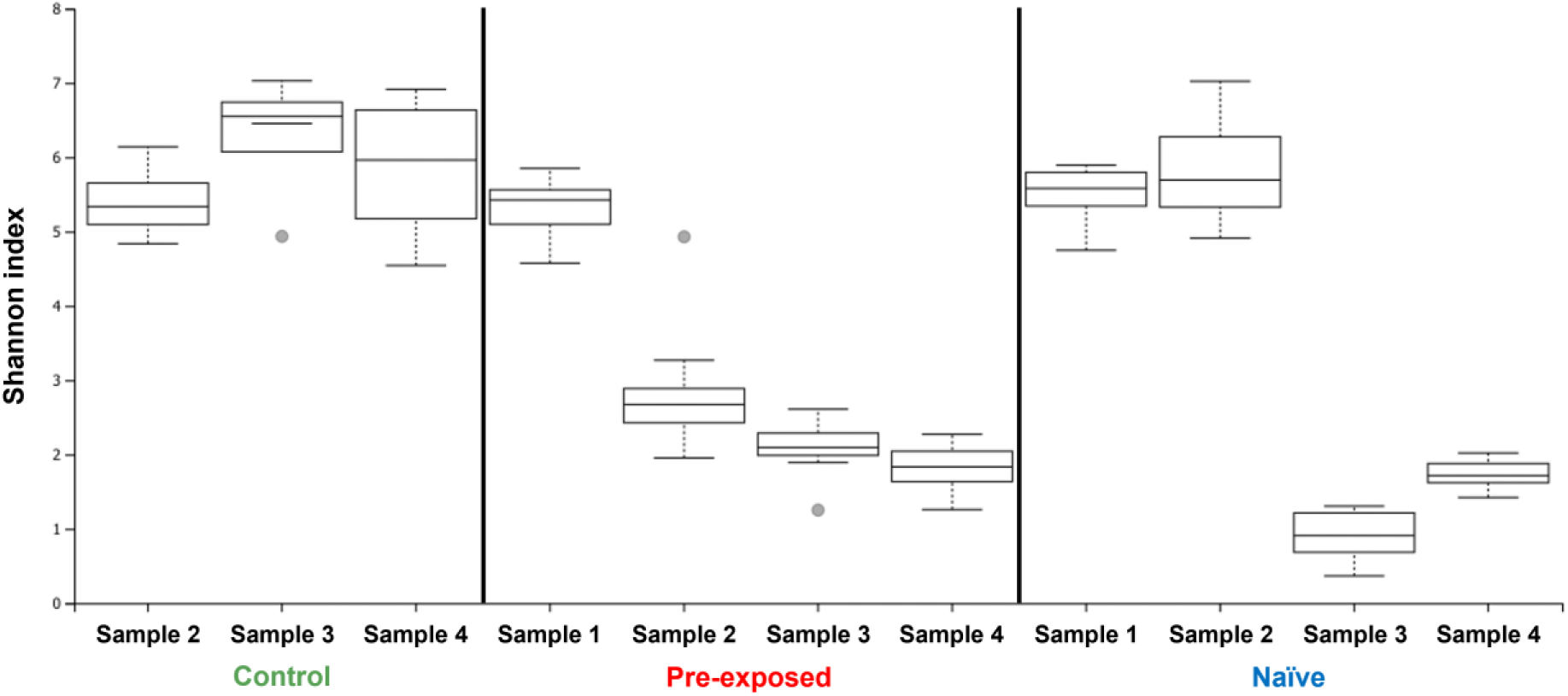
Shannon index values for control, pre-exposed, and naïve communities across time points. Significant differences were observed in pre-exposed communities between the start of the experiment (Sample 1) and end of phase I (Sample 2) (H’ = 18.64, *Q* = 0.000043) and between pre-exposed and naïve communities one week into phase II (Sample 3) (H’ = 22.91, *Q* = 0.000011). Pre-exposed and naïve communities converged by the end of the experiment (Sample 4) (H’ = 1.37, *Q* = 0.28). Control communities remained unchanged throughout the experiment (Sample 2 – Sample 3: H’ = 2.08, *Q* = 0.19; Sample 3 – Sample 4: H’ = 0.33, *Q* = 0.60). Benjamini-Hochberg *Q*-values based on *P*-values from Kruskal-Wallis tests.

**Figure S2.**
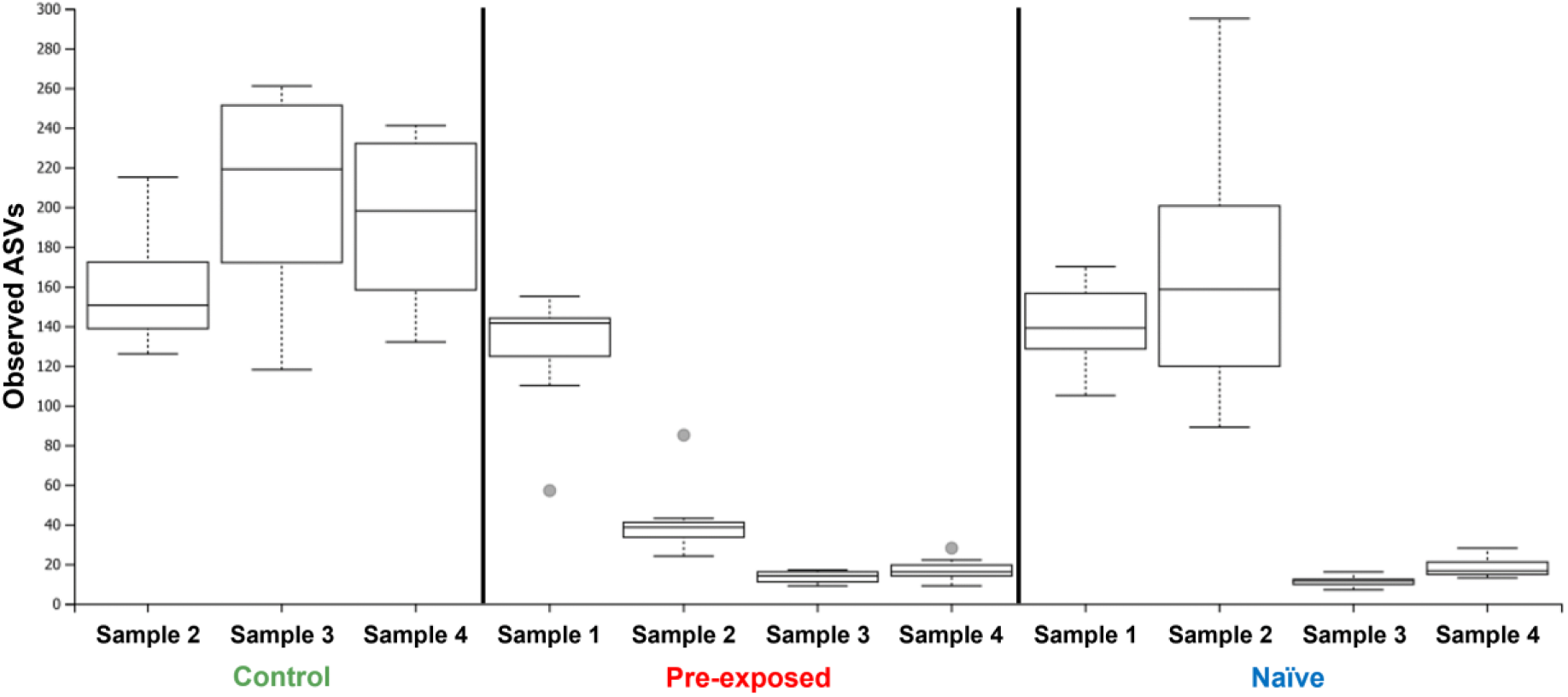
Observed number of amplicon sequence variants (ASVs) in control, pre-exposed, and naïve communities across time points. Observed ASVs were significantly greater in naïve communities at the end of the experiment (Sample 4) than immediately after severe acidification (Sample 3) (H’ = 17.95, *Q* = 0.000062) whereas no significant change was observed in pre-exposed communities (H’ = 4.11, *Q* = 0.060). Benjamini-Hochberg *Q*-values based on *P*-values from Kruskal-Wallis tests.

**Figure S3.**
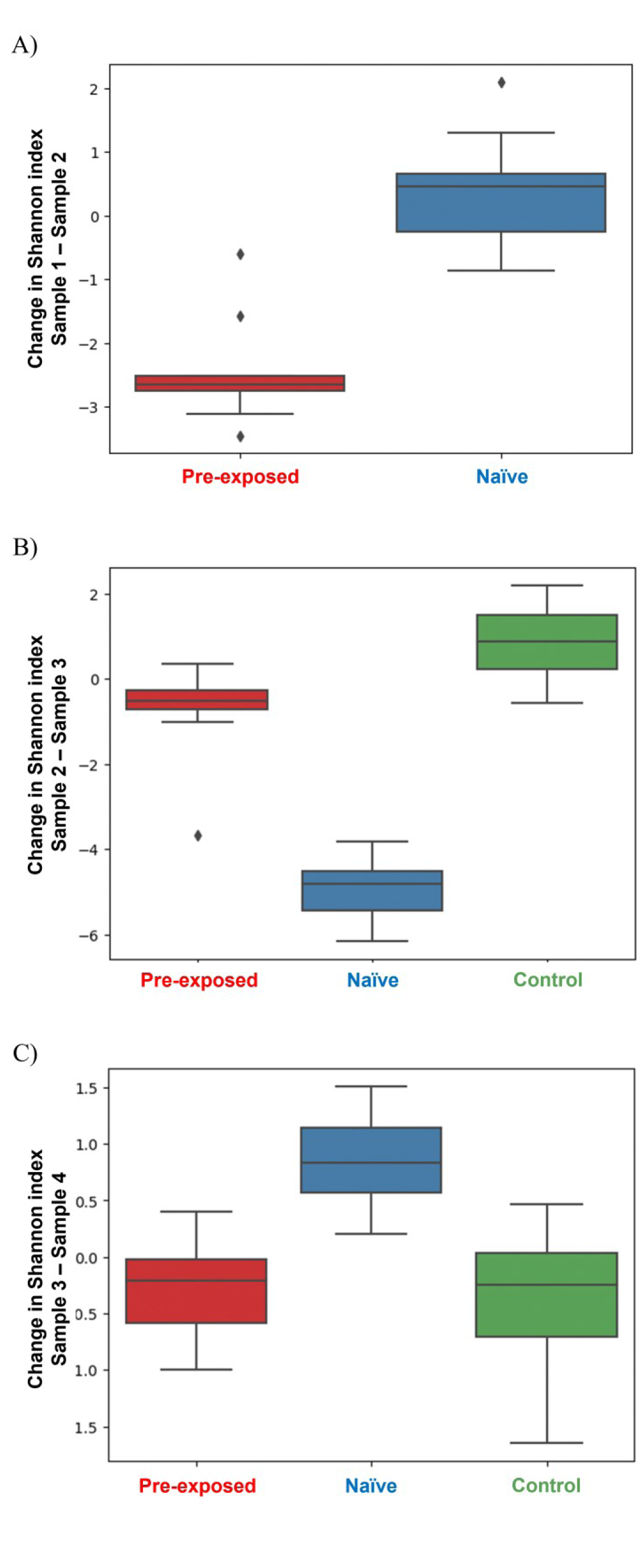
Pairwise change in Shannon diversity of pre-exposed, naïve, and control communities across time points. Significant differences were observed between pre-exposed and naïve communities from the A) start of the experiment to end of phase I (from Sample 1 to Sample 2) (U = 2.0, *Pfdr* = 0.00002), B) end of phase I to one week into phase II (from Sample 2 to Sample 3) (U = 240.0, *Pfdr* = 0.000007), and C) one week into phase II and the end of the experiment (from Sample 3 to Sample 4) (U = 2.0, *Pfdr* = 0.00001, Bonferroni *Pfdr*-values based on *P*-values from Mann-Whitney U tests). Pairwise change in the Shannon index of pre-exposed communities was significantly negative between all time points (Sample 1 – Sample 2: W = 0.0, *Pfdr* = 0.00098; Sample 2 – Sample 3: W = 6.0, *Pfdr* = 0.0013; Sample 3 – Sample 4: W = 20.0, *P_FDR_* = 0.032, Bonferroni *P_FDR_*-values based on *P*-values from Wilcoxon signed-rank tests) whereas it was only significantly negative in naïve communities immediately after severe acidification (Sample 2 – Sample 3: W = 0.0, *P_FDR_* = 0.000092) and significantly positive by the end of the experiment (Sample 3 – Sample 4: W = 0.0, *P_FDR_* = 0.000092), indicating recovery.

**Figure S4.**
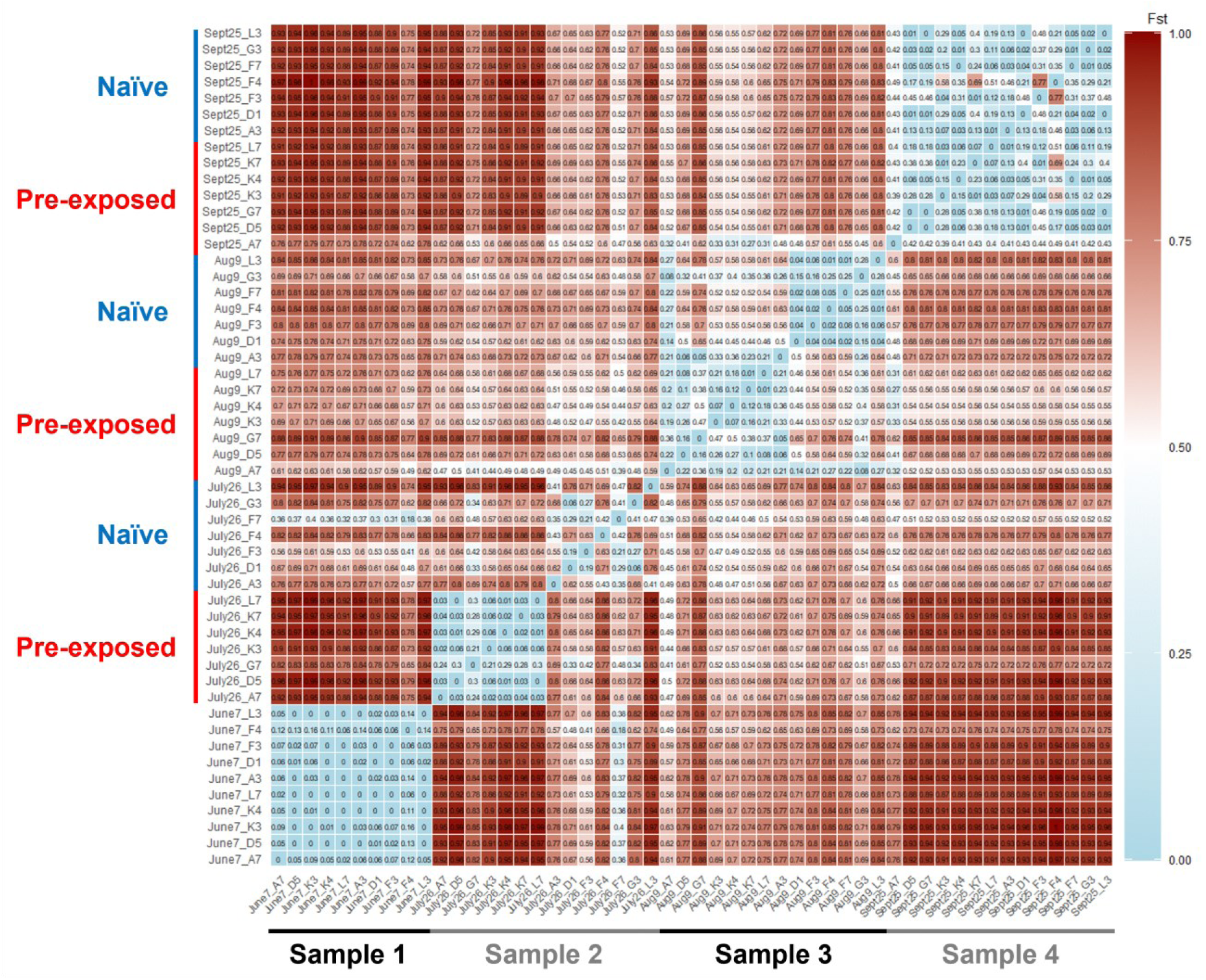
Pairwise fixation index (*F_ST_*) of *Acidiphilium rubrum* between each sample. Samples are ordered chronologically from left to right and from bottom to top. Time points are marked horizontally as Sample 1, Sample 2, Sample 3, and Sample 4. Phase I treatments are marked vertically as pre-exposed (red) and naïve (blue). *F_ST_* values ranged from 0 (light blue) to 1 (dark red).

**Figure S5.**
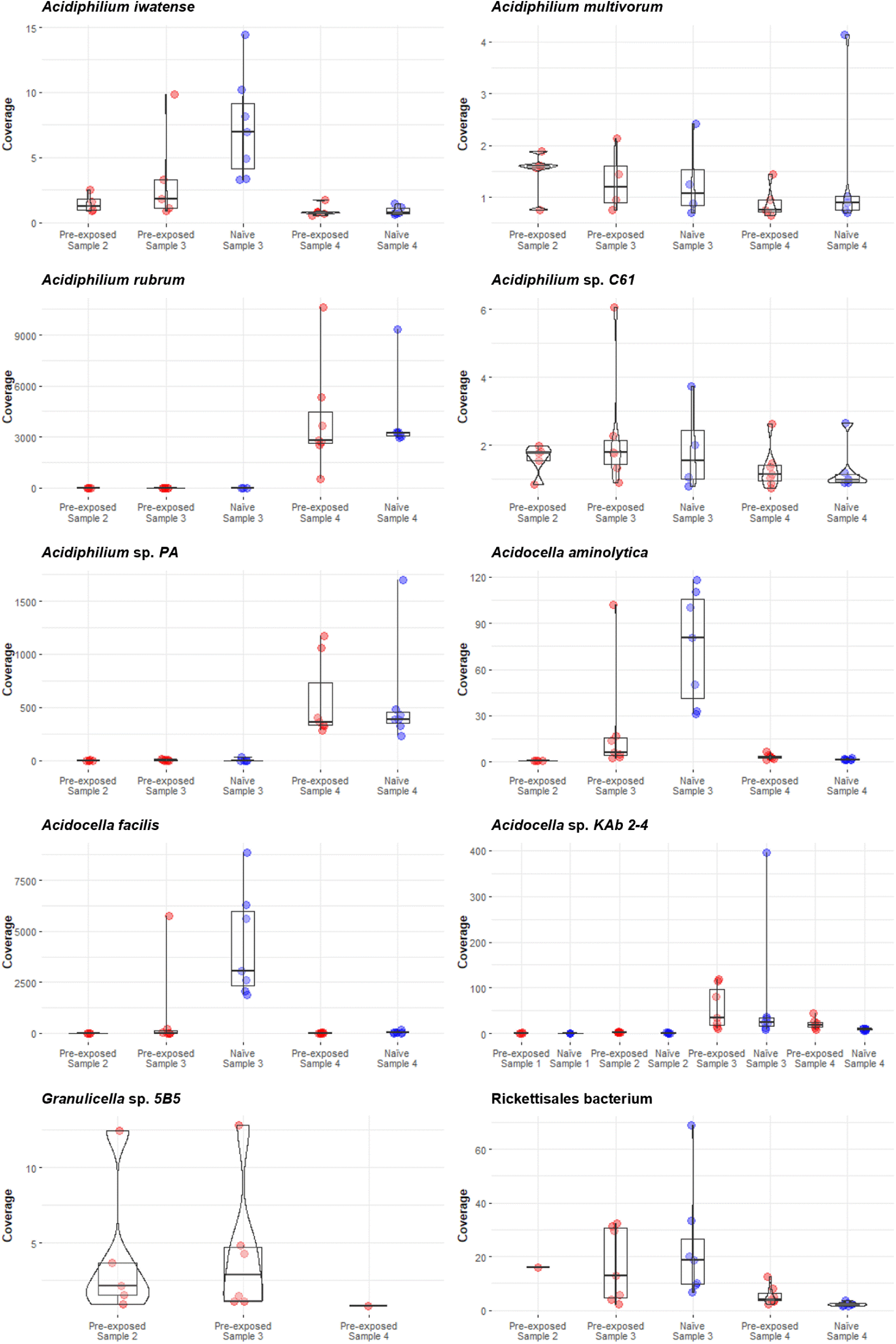
Mean coverage of genomes across species. Significant variation is observed between pre-exposed (red) and naïve (blue) communities across time. Each dot represents a distinct population from separate mesocosms.

**Figure S6.**
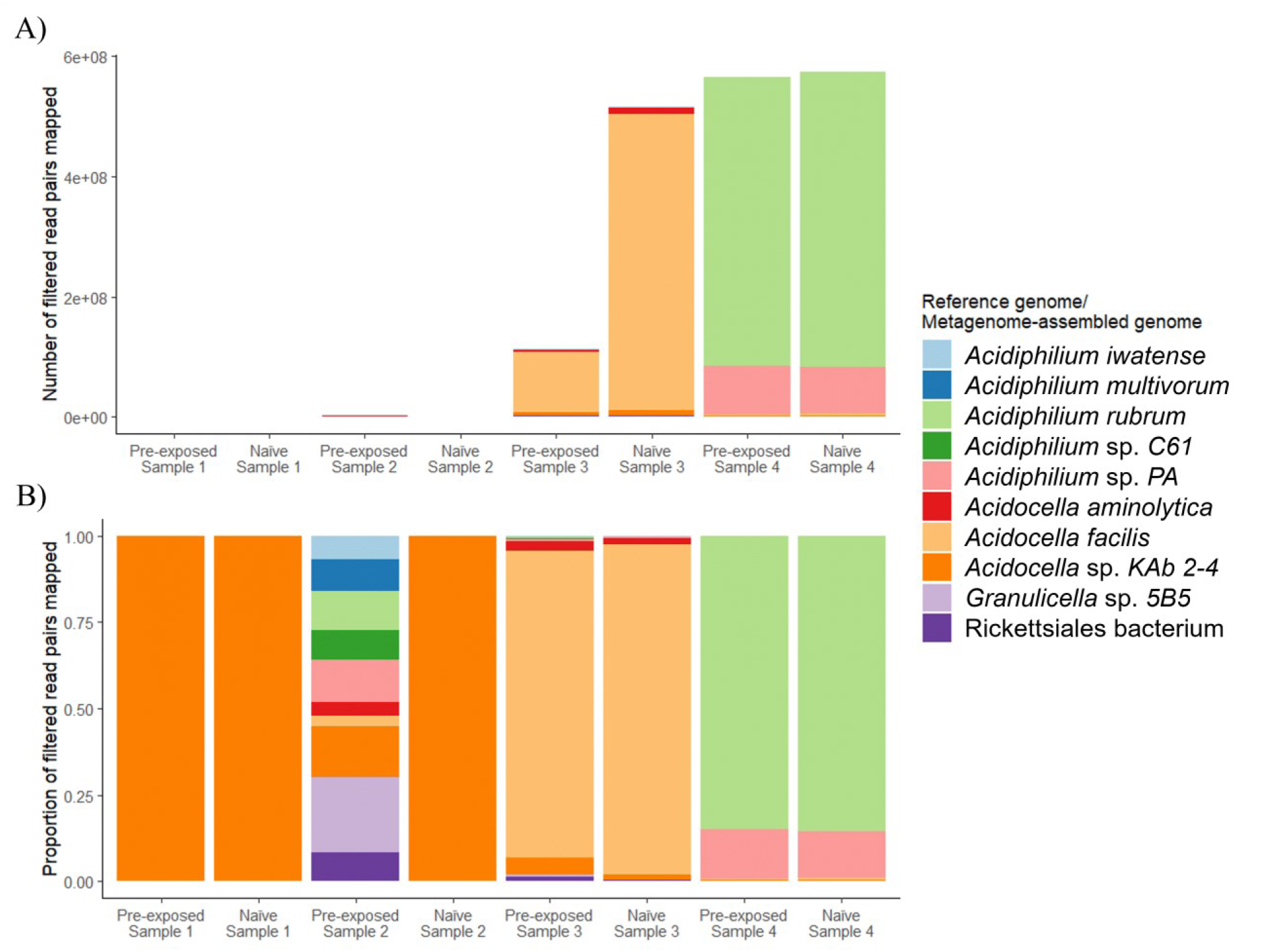
Read pairs mapped to reference genomes and metagenome-assembled genomes (MAGs) A) Total read pairs mapped. B) Proportion of reads pairs mapped.

**Figure S7.**
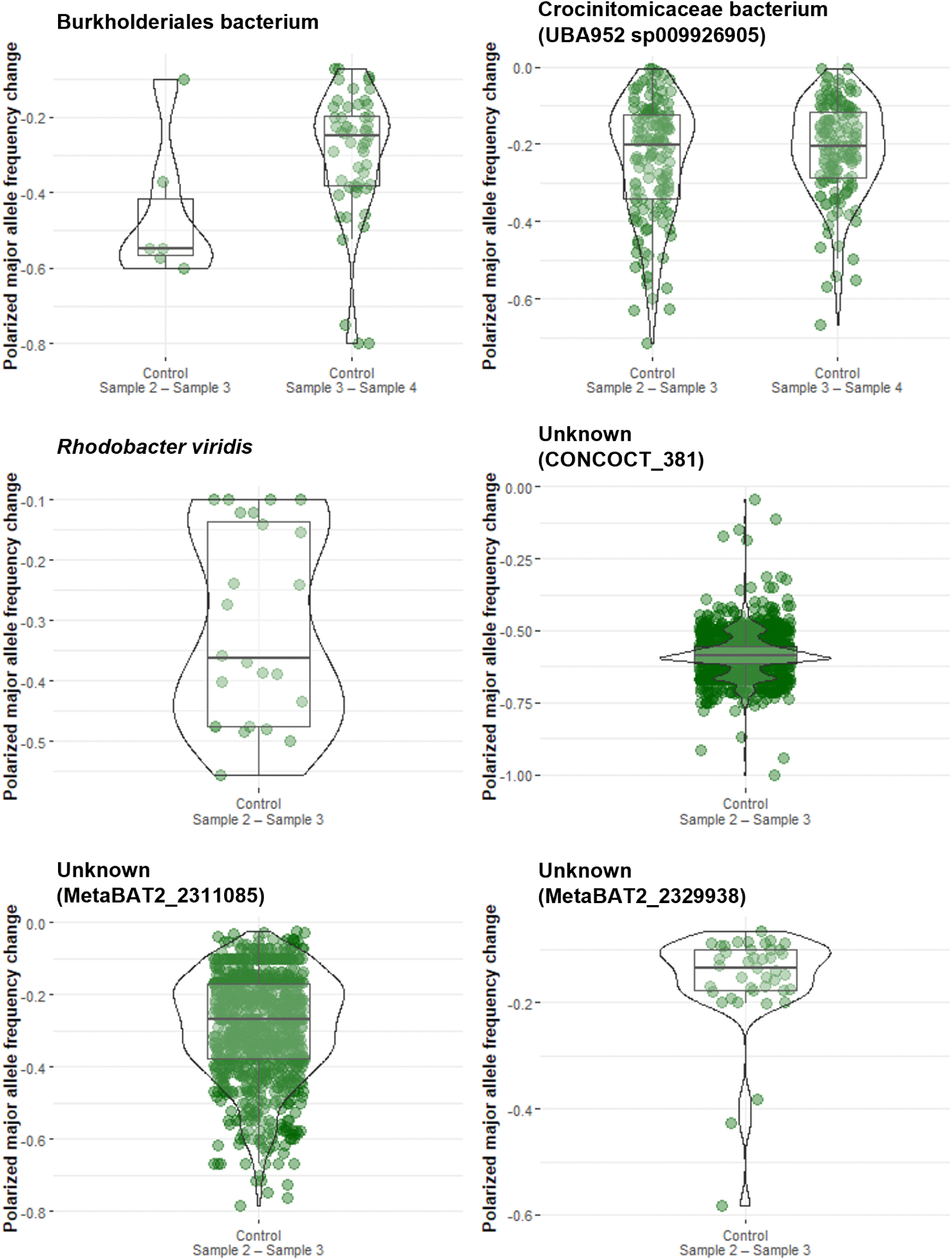
Allele frequency changes in single nucleotide polymorphisms (SNPs) polarized to major alleles of metagenome-assembled genomes (MAGs) in control communities. We calculated allele frequency change by subtracting the frequency of the major allele by the frequency of that same allele in a subsequent time point. Unnamed genus and species are in parentheses as well as MAG names if no taxonomic identity was able to be assigned.

## Notes

**Declaration of interests:** The authors declare no competing interest.

### Competing Interest Statement

The authors have declared no competing interest.

